# An Empirical Analysis of Topic Modeling for Mining Cancer Clinical Notes

**DOI:** 10.1101/062307

**Authors:** Katherine Redfield Chang, Xinghua Lou, Theofanis Karaletsos, Christopher Crosbie, Stuart Gardos, David Artz, Gunnar Rätsch

## Abstract

Using a variety of techniques including Topic Modeling, Principal Component Analysis and Bi-clustering, we explore electronic patient records in the form of unstructured clinical notes and genetic mutation test results. Our ultimate goal is to gain insight into a unique body of clinical data, specifically regarding the topics discussed within the note content and relationships between patient clinical notes and their underlying genetics.

## I. INTRODUCTION

Unstructured medical text notes contain a variety of information regarding patients and their care. Text data includes details such as family history, physician’s care plans, and current symptoms. Much of this information cannot be found anywhere else in a patient’s Electronic Health Record (EHR). By employing generative topic models tailored to this source of rich patient data we can gain a deeper understanding of these patients. Analyzing the correlations between patient clinical text and genetic testing results could reveal unexpected patterns.

In this work, we describe our exploration of a largely untapped set of EHRs containing patients’ initial consult clinical notes and the genetic mutations in those patients’ tumors. We begin by using topic modeling to characterize the entire corpus of notes. Then we apply principal component analysis to further reduce the dimensionality of patient note content, and use the first two principal components to search for possible interesting clusters amongst patients. Finally, we attempt to uncover correlations between patient notes and gene mutations in those patient’s tumors.

## II. RELATED WORK

The recent note of [1] is most similar to our work. There, various dimensionality reduction techniques were used to obtain a latent representation of patient state from clinical text analysis in an Intensive Care Unit setting. The study showed that vanilla unsupervised latent dirichlet allocation (LDA) [2] outperforms its supervised counterparts sLDA [3] and MedLDA [4] in a high-dimensional setting, which is the common case when attempting to represent the complexities inherent in patient data. This motivates our usage of vanilla LDA with more than 50 topics. In their work the focus was on studying predictability of clinical events such as sepsis based on the latent representations, while our setting is more exploratory, i.e. visualizing the data to find trends and studying correlations between genetic variables and latent patient states. Another difference may be that we went to great lengths to preprocess the notes (see Section IV-A), which they also motivate as a future need to build better representations.

In other related work, the promise of topic models to create abstractions from medication combination was studied [5]. Similarly, the interactions between herbs used in traditional Chinese medicine for given symptoms were analyzed in [6] using variants of LDA. However, clinical text mining using topic models is a crowded field, already. In other work [7], topic model representations of unstructured clinical text are used to classify and represent radiology reports. In a more general setting, biomedical text has been probed for relations between headings and content in using topic models and adaptations like Topic-Concept models in [8]. Other applications of similar models have been the exploration of recreational drug discussions [9] and, relevant to clinical practice, clinical case retrieval [10]. One key difference of our work to all of the above is the focus on cancer patient notes and the association to biological variables like genetics.

## III. DATA DESCRIPTION

### A. *The Darwin Database*

Darwin is a self-service reporting and analytics tool developed and maintained by Memorial Sloan-Kettering Cancer Center (MSKCC) which contains information on over 900 labeled data fields across 30 data subject areas. Over 1.3 million unique patients records have been added to date with information spanning back three decades. Darwin was created to help meet the increasing data demands of MSKCC’s research and clinical staff (see Figure 1).

**Fig. 1.**
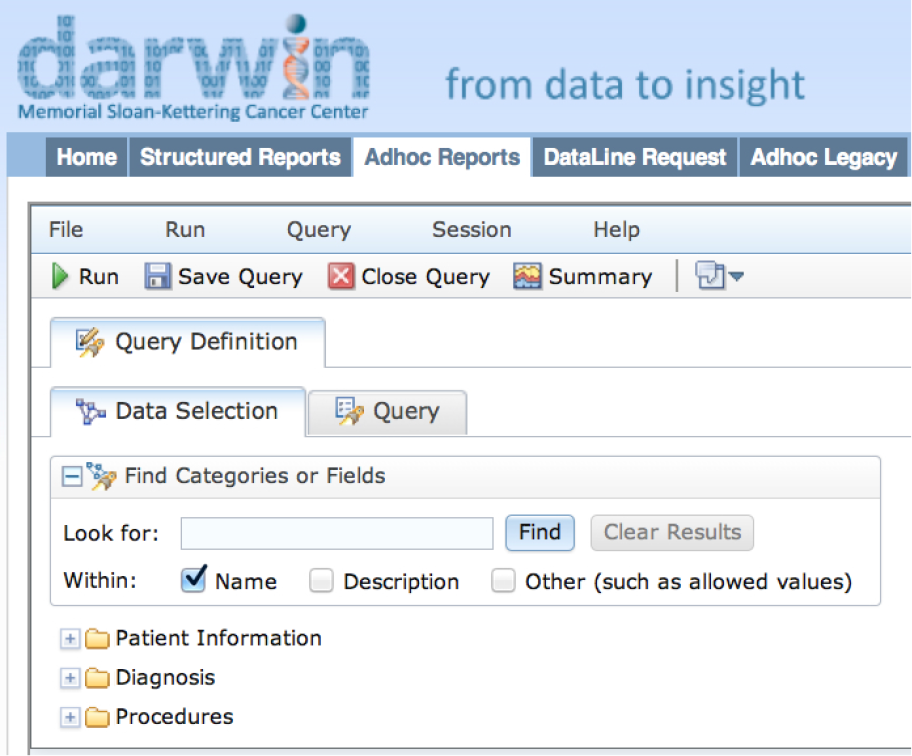
The user interface of the Darwin reporting system.

### B. *Data Selection*

Due to the Health Insurance portability and Accountability Act (HIpAA), patient data is strictly regulated. HIpAA’s “minimum necessary standard” requires medical entities to limit the disclosure of protected health information [11]. In order to comply with this regulation we selected a subset of patients and data fields from the EHR database to focus on in this first exploration.

Our dataset consists of 5,605 de-identified patient records for whom we could access both initial consult notes, and at least one genetic mutation test result from a common testing panel named Sequenom [12]. Since we wished to examine possible correlations between patient text and mutations, we chose these patients specifically because of the existence of their Sequenom gene mutation results.

The clinical notes are in the form of unstructured free text, and the Sequenom results are formatted as columns of discrete mutation “Positive” or “Negative” test results for each patient record.

## IV. METHODOLOGY

### A. *Data Preparation*

Data pre-processing is an essential step before applying any downstream data mining or machine learning techniques. Irrelevant, noisy and unreliable data makes knowledge discovery more difficult and increases the likelihood of producing misleading results [13].

In order to improve the quality of our analysis, we took several steps to prepare our raw patient data into a more consistent and regular dataset. For the medical text we removed stopwords, applied negation detection, stemmed each term, and separated the content into logical subsections before starting our topic modeling. For the genetic testing results, we discarded sparsely documented mutations, patients with contradictory results, and duplicate patient entries.

**Fig. 2.**
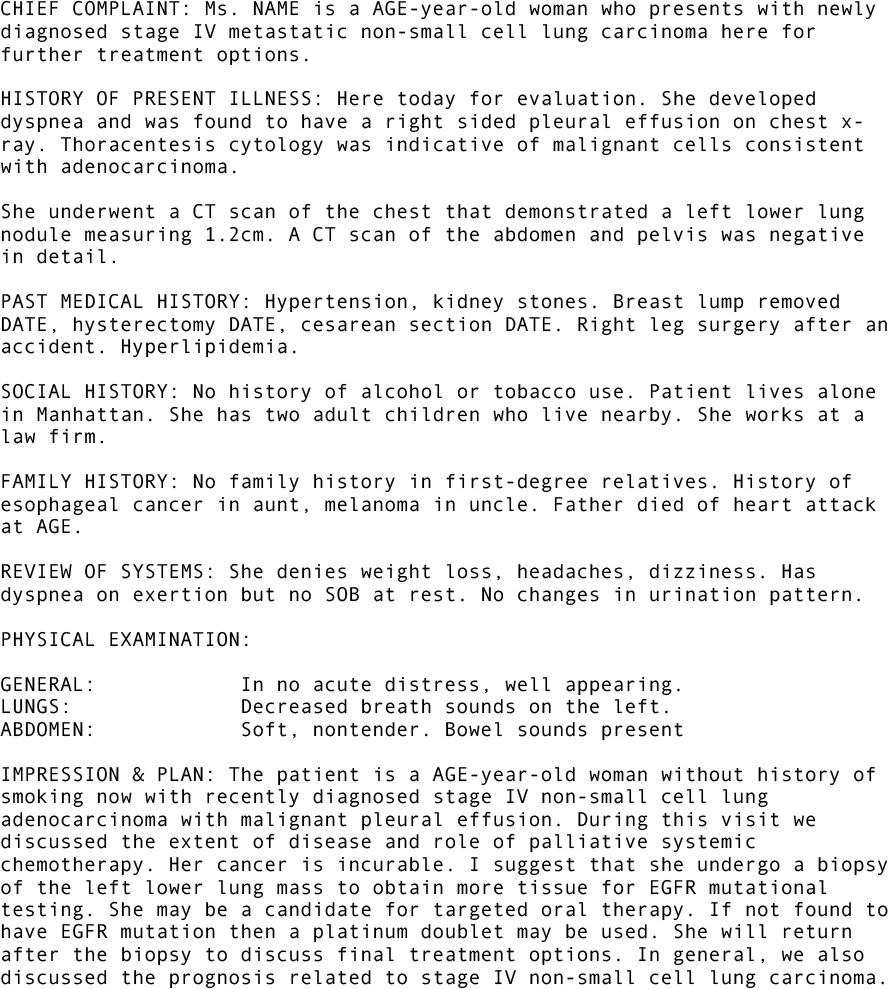
A Sample Clinical Note

#### 1) *Clinical Text Preparation:*

First we removed extremely common stopwords with little contextual value such as ‘the’, ‘and’, and ‘it’. We started compiling our list of stopwords from an existing list of common English stopwords [^1^ http://www.ranks.nl/resources/stopwords.html] and added in new ones that were more specific to our dataset such as keywords for redacted information like ‘DATE’, ‘INSTITUTION’ and ‘NAME’.

Next we performed negation detection on our text because the presence or absence of symptoms and behaviors is one of the most fundamental pieces of information available in clinical notes. To capture this concept we first replaced verbose negative phrases with the word “no” to reduce variability in the data. Then, if a negating phrase or word appeared at the beginning of a list it was applied to the entire list. Lastly, in order to retain the negated state of a given word, we linked negative identifiers to the words they negated to be treated as their own distinct terms. For example, “no history of tobacco, drugs, alcohol” becomes “no_tobacco no_drugs no_alcohol”.

In order to avoid duplication of concepts, we chose to treat similar words as one concept. For example, “compute”, “computing”, “computes”, and “computed” would all be treated as instances of one word group with the stem “comput”. To make the resulting text easier to read, when building our document vocabulary we treated all instances of a particular word-stem as instances of the first word identified by that stem. We implemented this stemming process using the python stemming package’s Porter2 algorithm [^2^ https://pypi.python.org/pypi/stemming/1.0].

**Fig. 3.**
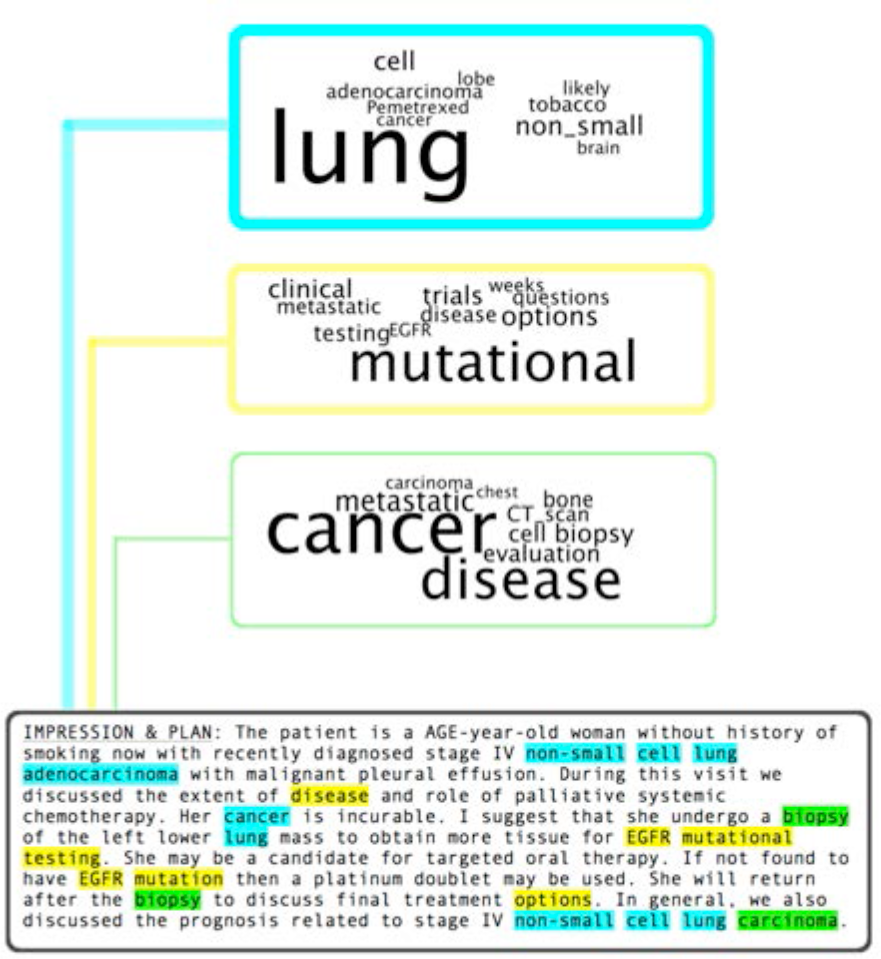
Sample Topic Model Results – the word clouds represent topics which were generated over the entire body of documents, the highlighted words in the text show which words from this note are attributed to the matching colored topic. Line thickness indicates the percentage of text attributed to that topic.

Clinical note content varies dramatically from physician to physician, institution to institution, and sometimes just day to day. This problem is further compounded by the fact that medical terminology is not standardized. There are a plethora of synonyms, acronyms and abbreviations as well as spelling errors. While we considered using a medical dictionary like the Unified Medical Language System (UMLS) [14], to standardize the document language, we eventually abandoned that idea as often abbreviations and acronyms stand for multiple different concepts, and because we did not want to lose detail by treating more specific terms as instances of their parent concept (for example, “adenocarcinoma” contains more contextual information than “carcinoma”).

One of the only consistencies in note content across different patients and physicians was the basic outline. Commonly, clinical notes are composed of several distinct sections: Chief Complaint (CC), History of Present Illness (HPI), Past Medical History (PMH), Family History (FHx), Social History (SHx), Review of Systems and Physical (ROS) and the physician’s Impressions and Plan (IMP) (see Figure 2). Since different sections focus on different information and share contextual similarities from one patient to the next, we decided to examine each section individually. We corrected for common spelling errors and variations in labeling these sections, and then used a rule-based system to divide the notes into these components.

**Fig. 4.**
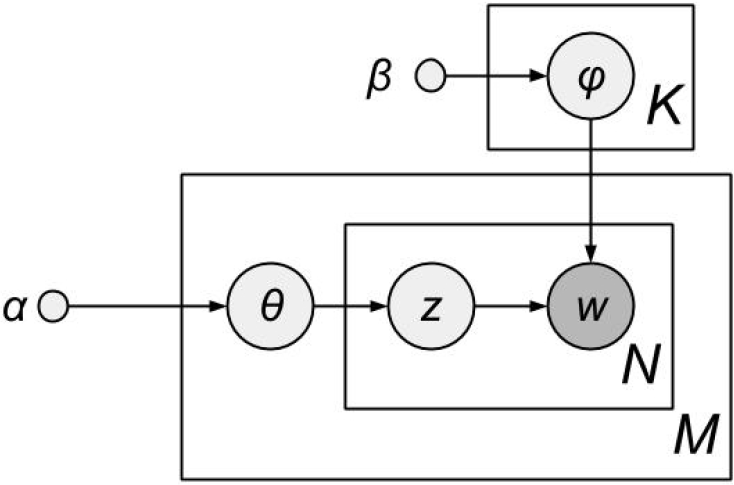
Graphical Model Representation of LDA

#### 2) *Sequenom Preparation:*

Each patient had at least one genetic test result, though not all patients had the entire panel’s results loaded into the EHR database. To reduce the likelihood of generating biased results, we chose to examine those genetic mutations for which at least 1000 patient results existed, and which identified 5 or more patients as “positive”. We removed from consideration patients with contradictory records (both positive and negative test results recorded). Last, duplicate entries were removed.

### B. *Topic Modeling*

Topic models are a class of statistical models used in natural language processing to reveal the underlying thematic structure of a large body of documents [2]. The intuition behind topic modeling algorithms is that a document discussing a certain concept is more likely to contain language associated with that concept. Intuitively, if a document is 80% about lung cancer and 20% about alternative medicine, then approximately 80% of the terms in that document will be related to lung cancer and approximately 20% of the terms will be related to alternative medicine (see Figure 3). A topic is simply a distribution of terms over a vocabulary, allowing each document to be described as a distribution over topics.

One of the most commonly used unsupervised topic modeling algorithms is latent Dirichlet allocation (LDA) [15]. LDA resembles a generative process that assembles a document collection ***D*** = {d_*m*_}_*m*∈{1…*M*}_ using a fixed vocabulary ***W***, subject to some unknown probability distributions including the distribution of topic *k* over vocabulary (denoted as **Φ** = {ϕ*^k^*}, *k* ∈ {1,…, *K*}) and the distribution of the mth document over all *K* topics (denoted as **Θ** = {*θ_m_*}, *m* ∈ {1,…, *m*}). LDA can be represented as a probabilistic graphical model in Figure 4, where *z* is a topic assignment vector for words *w*, and *α* and *β* are prior parameters. This model also corresponds to Eq. (1) which clearly describes the generative process: for document *m*, first the distribution of topics over vocabulary **Φ** and the distribution of document over topics *θ_m_* are sampled from prior *β* and *α*, respectively; then the topic assignment *z* for each word is generated from *θ_m_*; and finally the exact words *w* are generated according to their respective topic assignment *z* as well as the distribution of topics over the vocabulary **Φ**.

**Fig. 5.**
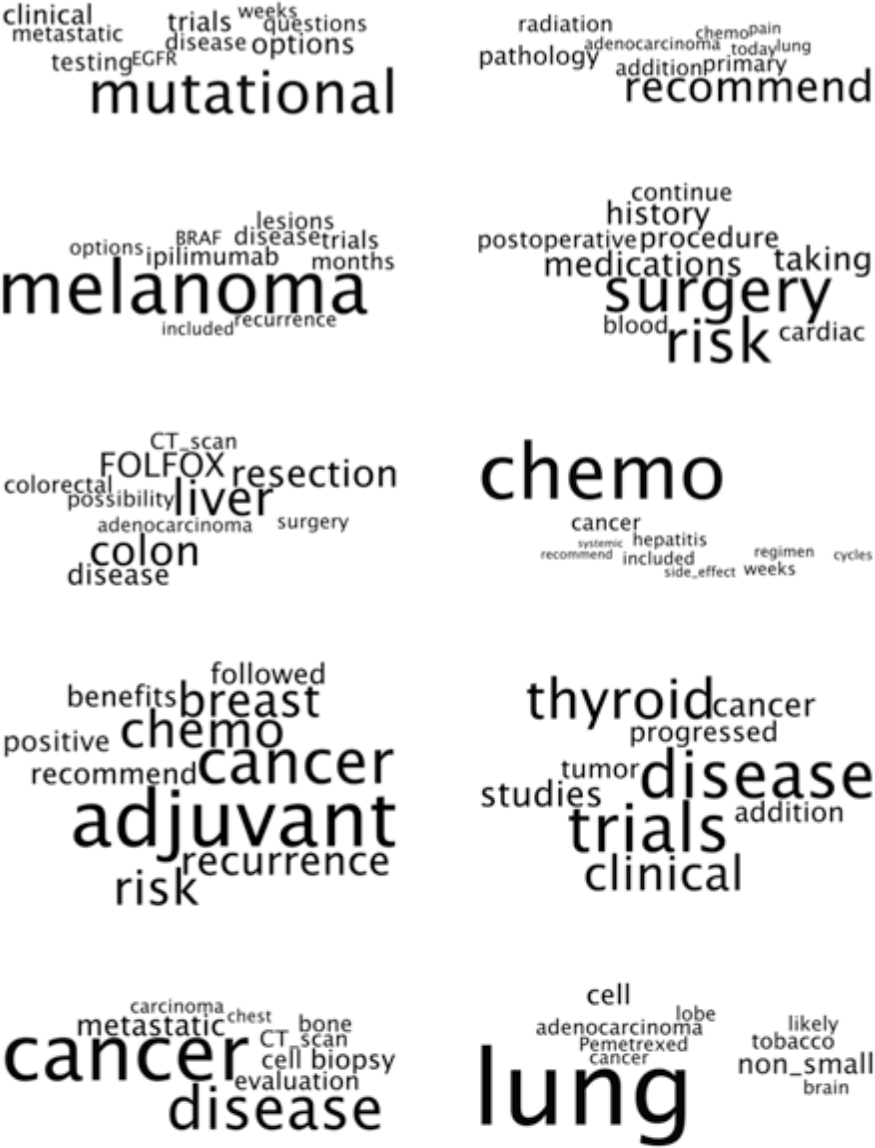
Topic modeling output with ten topics for the Impressions & Plan section of this corpus of patient notes. Each word cloud represents one topic discovered using LDA. A topic is a distribution over words; here, relative word size shows the relative weight of a given word within the topic distribution.

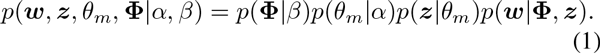

To extract **Φ** and **Θ** that are of our interest, we can use maximum log-likelihood estimation over ***D***. Unfortunately, the discrete variable *z* causes this inference to be intractable in practice. This issue has been solved using collapsed Gibbs sampling [16], an algorithm for inference in LDA. Specifically, we used LDA with collapsed Gibbs sampling as implemented in the GraphLab Topic Modeling toolkit [17] for our experiments.

## C. *Principal Component Analysis and Clustering*

In principal component analysis (PCA), a dataset is orthogonally transformed into linearly uncorrelated bases. These new bases, referred to as ‘principal components’ are defined such that the first principal component describes as much of the variability within the data as possible. Each following principal component is defined to be orthogonal to the preceding components to explain the maximum possible remaining variance.

**Fig. 6.**
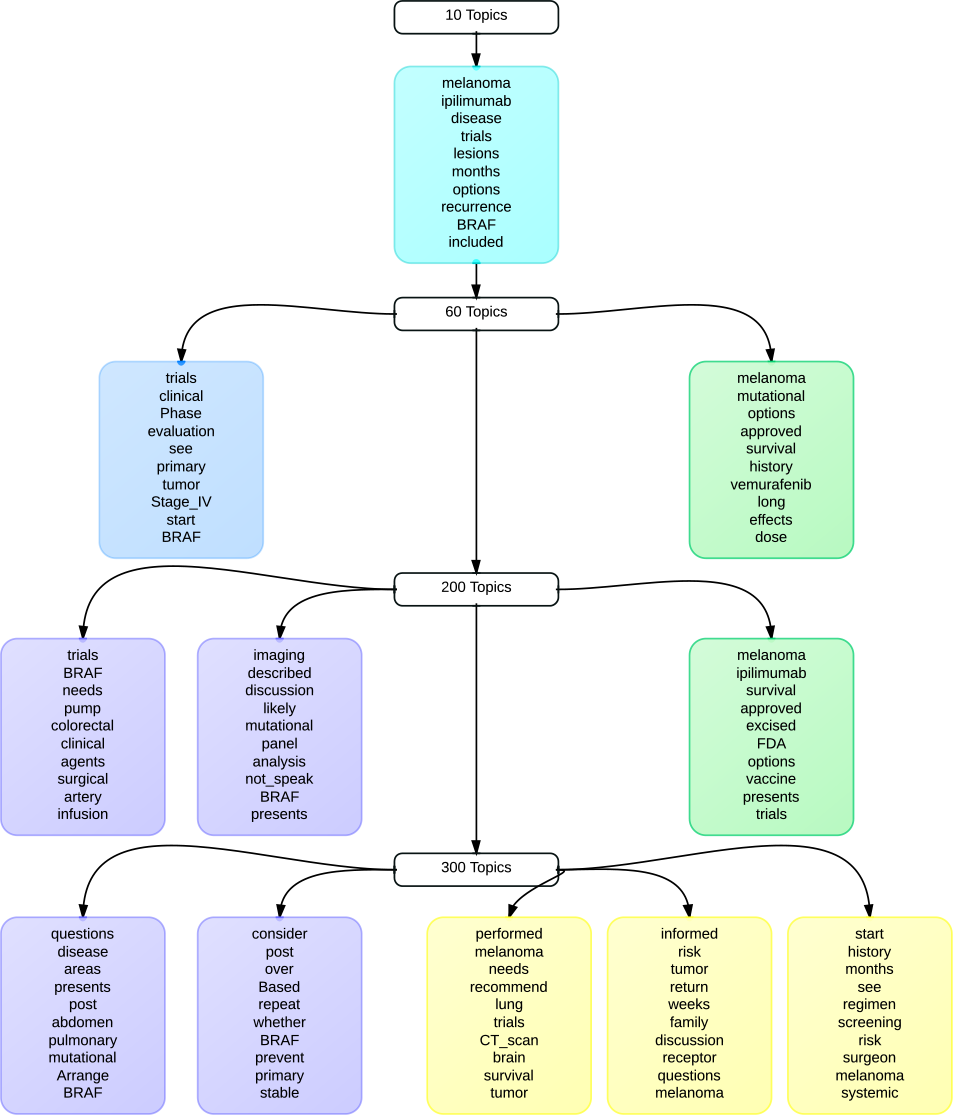
Topic Stability: Topics concerning ‘melanoma’ and ‘BRAF’ appear in every LDA output for the Impressions & Plan note section. These topics grow from a single broad category when LDA produces 10 Topics to more detailed word groups as more topics are computed.

Our goal in applying PCA to the topic distributions, **Θ**, was to reduce the dimensionality of our data sufficiently to be able to visualize all the patients simultaneously, and to see if patients could be grouped in some logical manner this way. We accomplished this aim by calculating and plotting the first two principal components for each section and topics group, and examining those plots for clusters using a meanshift clustering algorithm [18].

## D. *Mutation Correlation and Bi-clustering*

We selected this group of patient’s clinical notes from the EHR database specifically because these patients all had genetic mutation test results. One goal of this exploration was to see what relationships, if any, exist between the text of a patient’s notes and their subsequent genetic testing results. In order to explore these possible relationships, we examined each set of mutation results as they related to each topic in several ways. First, for every combination of mutation and topic, we computed the Pearson’s correlation coefficient between the patients’ mutation results and the percentage of their notes attributed to the given topic. We then ranked these coefficients by their absolute value from largest to smallest, and compared the results of the strongest correlations. Finally, we used the algorithm from [19] to discover and visualize possible bi-clusters within the most highly correlated topics and mutations.

**Fig. 7.**
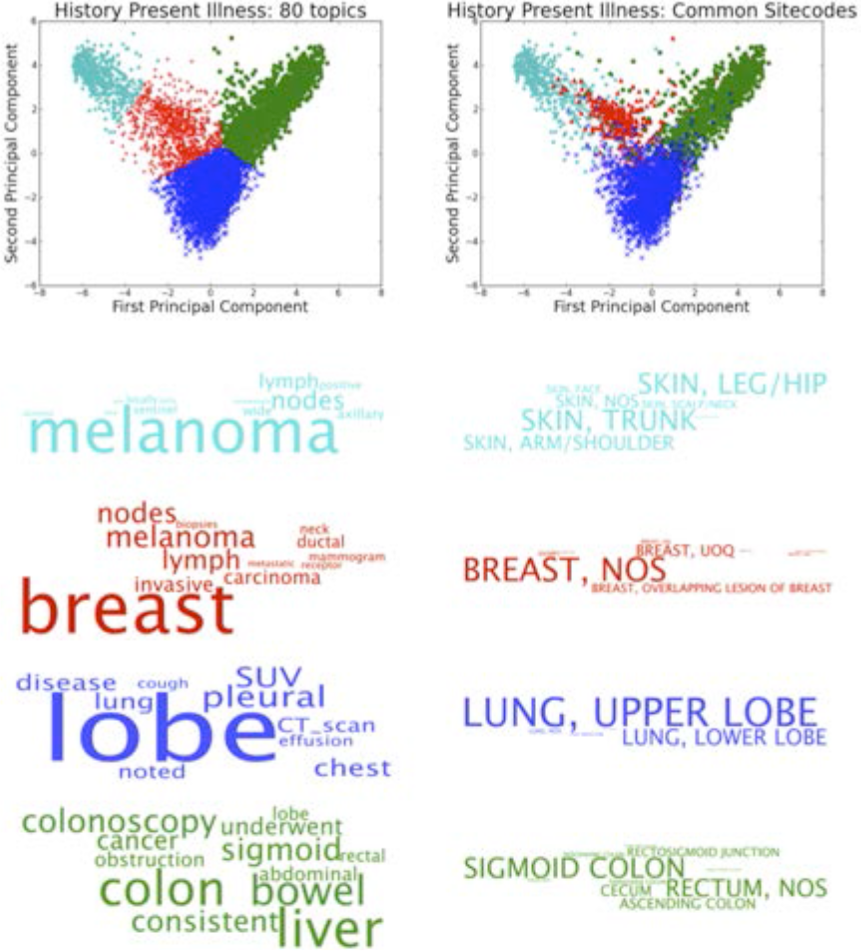
Left: The first two principal components, colored by meanshift clustering on History of Present Illness (HPI) for 80 Topics. Each cluster is described by the average topic distribution over patients. The resulting combination of topics is defined by a distribution of the top ten most common words from each component-topic. Right: Plot of the same first two principal components colored and defined by patient ICD site codes. Note the broad similarity of note-topic clusters, and groups seen by coloring those points via patient’s separately documented site code.

## V. RESULTS

### A. *Extracting and Visualizing Topics*

#### 1) *LDA for Topic extraction:*

We trained multi-scale topic models on each of the main note sections, generating groups of 10, 20, 30, 40, 50, 60, 70, 80, 90, 100, 200 and 300 topics for each section to provide an assortment of topic results for analysis. We then examined the resulting word clouds as a first-step sanity check to determine the quality of our extracted topics. The results of the LDA topic modeling appear to be descriptive, readable and reflective of the body of patients (Figure 5).

#### 2) *Topic Stability:*

While selecting different numbers of total topics generated a variety of results for analysis, we discovered that in most sections a few dominant topics would appear in every set of topic groups. This topic stability persisted from an initial broad category in the 10 topic range to more granular sub-categories on the 300 topics side (Figure 6).

#### 3) *Principal Component Analysis and Clustering:*

During billing for hospital services, cancer patients are assigned an International Classification of Diseases (ICD) Site Code describing the location of their tumors, infections and other injuries. In order to verify the clusters discovered via PCA, we decided to plot the patient points colored by their site codes. Since site code descriptions can be very specific (ex: SKIN, LEG/HIP) or relatively vague (ex: SKIN, NOS), we grouped together site codes by broad general location and show the most common results. Sometimes, the PCA clusters we discovered were descriptive of location-specific cancers such as Melanoma or Colon Cancer. As expected, we discovered that these topic clusters strongly resembles their site code plots (Figure 7).

**Fig. 8.**
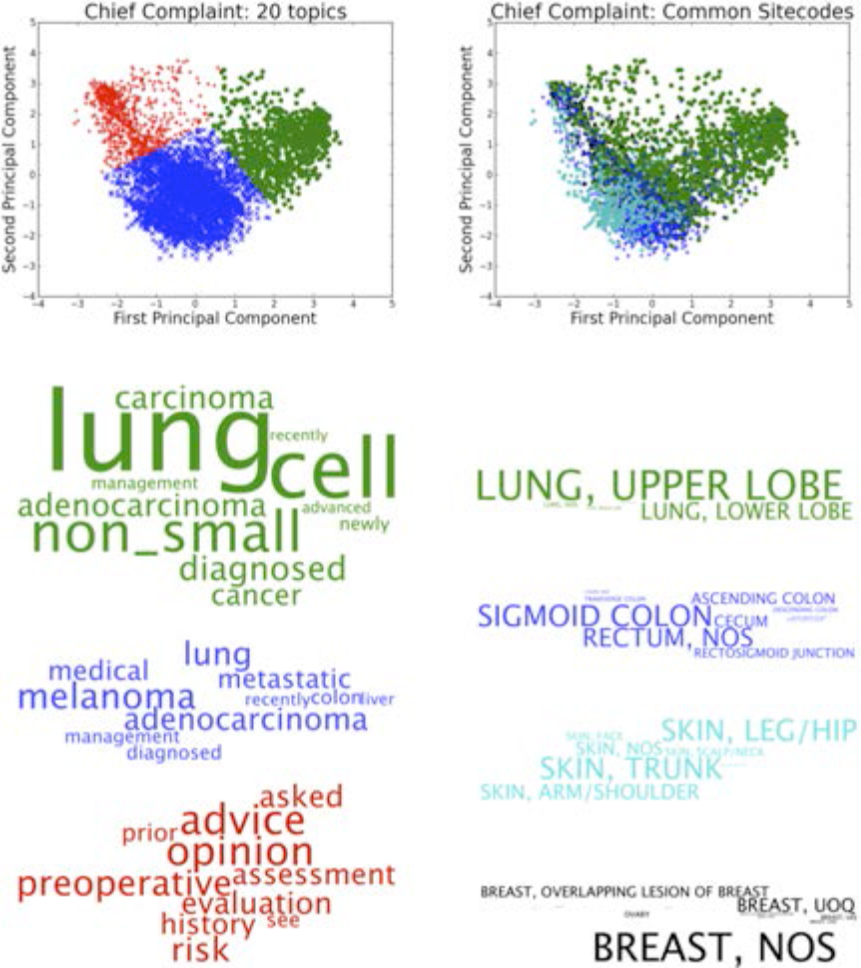
Chief Complaint Topic Clusters and Site Codes. Unlike Figure 7, here patient topic clusters do not seem to re-appear when patients are colored by site code. By examining the first two principal components of patient topic distributions we can reveal clusters of patient concern that are not immediately obvious from other metrics like billing site codes (such as preoperative risk assessment in red above).

Interestingly, principal component clusters of other sections described trends of patient concern that were less readily observable from raw patient statistics. For example, in the Chief Complaint section we discovered a large cluster of patients seeking advice and pre-operative risk evaluation. This cluster disappears in a site code plot (Figure 8).

### B. *Cross-Section Analysis*

As discussed above, clinical notes are composed of several sections. Each section describes a different aspect of a patient’s well-being and care. However, since each section is still discussing the same individual, some overlap in content is to be expected. In order to observe the level of similarity between sections, we computed correlations between patient topic distributions from section to section. In Figure 9 we highlight those topics with the highest correlations (*r* > 0.1 and *p* < 0.0005).

Clear visual patterns emerge showing high levels of similarity between History of Present Illness and Impressions and Plan, as well as slightly weaker but still visually striking relationships between these two sections with the Chief Complaint and Review of Systems/Physical. This is in contrast to Family History and Social History, which appear to lack significant correlation to other sectional content.

**Fig. 9.**
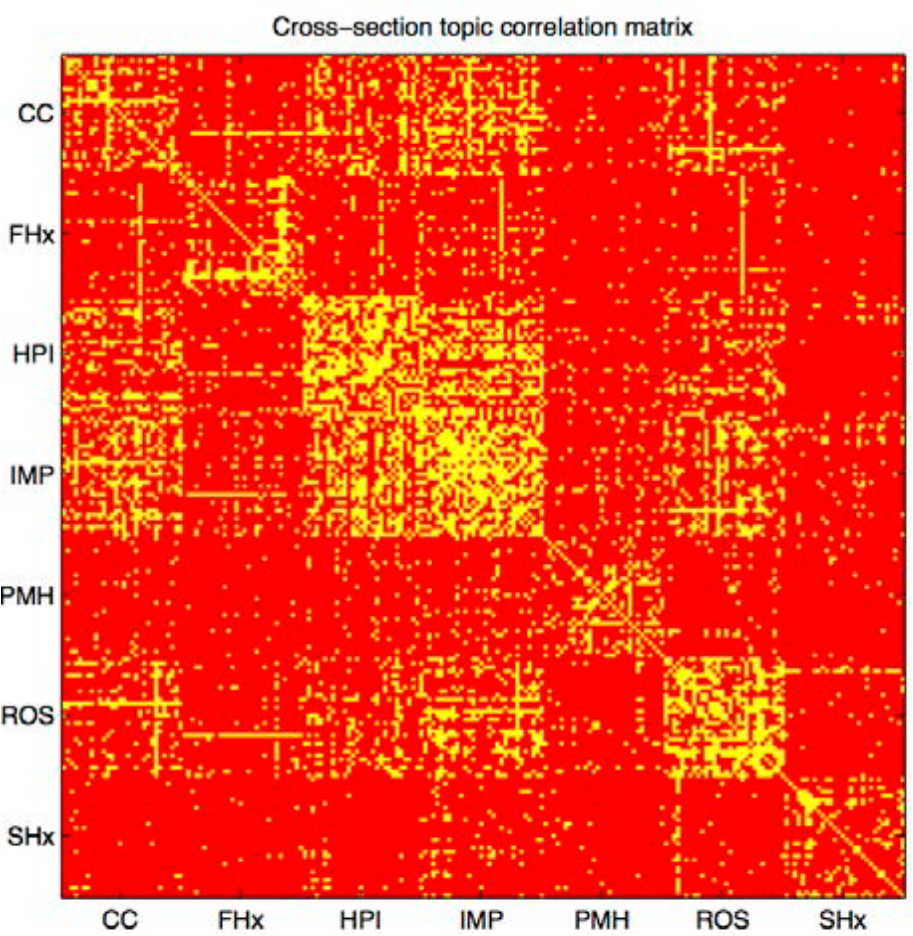
Cross-section topic correlation matrix. Topics with high Pearson correlation coefficient and low *p*-values are highlighted in yellow. Clusters appear to indicate significant overlap in content between HPI and IMP.

In context it seems sensible that a physician’s consult notes would show similarities between patient complaints, their present illness, bodily symptoms discussed during a physical review and the doctor’s impressions. Meanwhile, Family History and Social History contain many unique patient details. From this analysis it appears those details are topically unrelated to patient illness.

### C. *Mutation Correlation*

The Sequenom panel tests for mutations that are already known to exist in certain types of cancers. The goal of this correlation study was to see if we could independently re-identify any known relationships between mutations and cancer phenotypes as a proof-of-concept to test the reliability of using topic modeling to generate useful labeling of patients and find meaningful correlations. If we examine the strongest correlations between patient topics and mutation test results (those where *r* > 0.1 and *p* < 0.0005), we find several interesting correlations.

We discovered several topics with notable correlations to specific genetic mutations. First, we examined individually these strongest correlations between patient topics and mutations results. We found that 0.27% of topic-mutation pairs were notably correlated. In Table 1 we show the specific individual top correlations for the Impressions & Plan section with 20 topic groups. When reviewing the highest correlated items we repeatedly noted topics correlating to the following mutations: BRAF-V600, several EGFR mutations, KRAS-A146, NRAS-Q61, and PIK3CA-H1047. Furthermore, these mutations paired with multiple topics but because of the topic stability and overlap in sectional content discussed earlier, we were able to see a pattern of content emerge from these correlated topics. Table II shows the most common topic content associated with these mutations.

**Table I.**
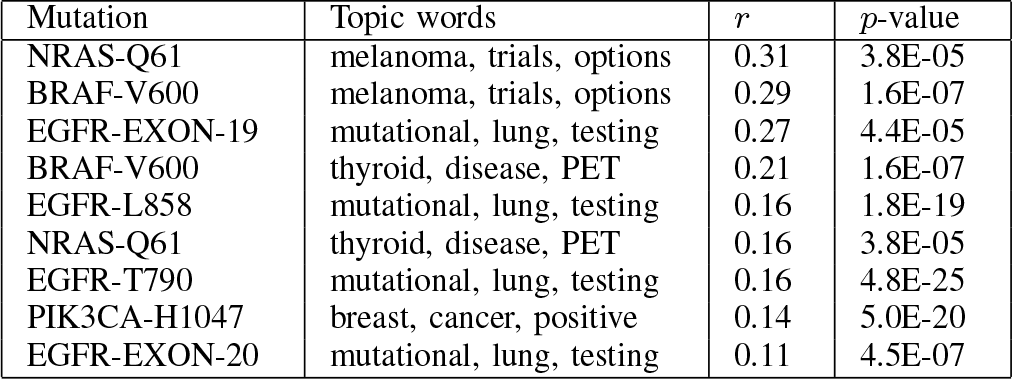
TOP MUTATION-TOPIC CORRELATIONS BETWEEN POSITIVE MUTATION TESTS AND IMPRESSIONS & PLAN 20 TOPIC GROUP.

**Table II.** REPEATED MUTATION-TOPIC CORRELATIONS

| Mutation | Correlated topic words |
| --- | --- |
| BRAF-V600 | melanoma, excision, malignant, wide |
| BRAF-V600 | thyroid, papillary, carcinoma, neck |
| BRAF-V600 | BRAF, melanoma, ipilimumab, clinical |
| EGFR-* | lung, adenocarcinoma, erlotinib, Stage_IV |
| EGFR-* | EGFR, mutation, lung, no_tobacco |
| KRAS-A146 | colon, metastatic, liver, sigmoid |
| NRAS-Q61 | melanoma, excision, malignant, wide |
| NRAS-Q61 | thyroid, papillary, carcinoma, neck |
| PIK3CA-H1047 | breast, positive, invasive, ductal |

We first notice that for BRAF-V600 and the EGFR mutations that ‘BRAF’ and ‘EGFR’ show up in commonly correlated topics. This is promising since it implies that our clustering and correlation study is finding reasonable correlations between note content and mutation results. More importantly, these correlations also find notable relationships in less obvious areas. For example, “NRAS” is never seen in the topic content correlating NRAS-Q61 to melanoma and thyroid cancer. For EGFR mutations, we see a relationship to lung cancer and erlotinib outside the topics containing “EGFR””.

In the bi-clustering step, we categorized each relationship between a topic and a mutation using the same standards for labeling the strongest correlations. Then, using the algorithm described in [19] we learn bi-clusters of correlated mutations and topics. Using this process we discover that BRAF-V600 and NRAS-Q61 mutations correlate to the same topics as each other, and can visualize the cluster between EGFR-EXON-19, EGFR-EX0N-20, EGFR-L858 and EGFR-T790 and topics containing “EGFR, mutation, no_tobacco” and “erlotinib, lung, adenocarcinoma.” See Fig. 10 for the biclustering results discovered between the Sequenom mutations and Chief Complaint 100 topics.

Since the Sequenom panel tests for mutations that arealready known to exist in certain types of cancers, we can examine the other correlated topics to determine if we were successful in independently identifying any verified relationships between mutations and cancer phenotypes.

**Fig. 10.**
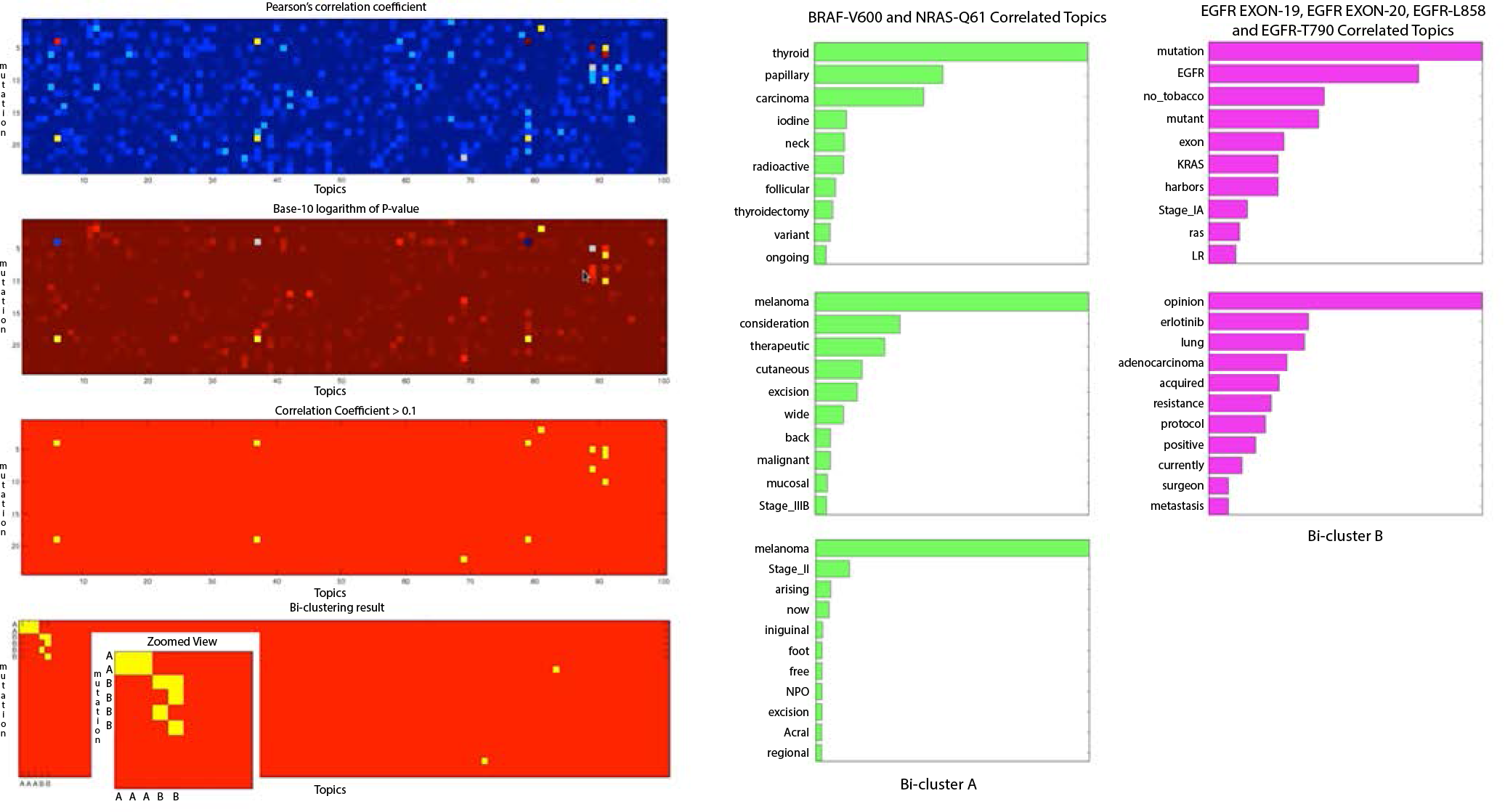
On the left the bi-clustering process is demonstrated. The blue heatmap shows the range of correlation between topics and mutations. The dark red heatmap shows the range of p-values for the same points. We combine these two datasets to identify clusters of the strongest correlations between topics and mutations. Those strongest relationships are highlighted in yellow in the third heatmap. The last step shows the bi-clustering results. The right-hand side shows the content of these bi-clusters. The green topics correlated to BRAF-V600 and NRAS-Q61 represent the top left-cluster with two mutations and three topics, and the magenta topics correlated to several EGFR mutations represent the second-cluster.

Excitingly, these strongest relationships have been detected independently by other methods of research [20], [21], [22], [23], [24]. This success in identifying independently verified relationships to mutations encourages us to speculate that refining and expanding upon this approach would be a valid avenue for future study with a larger corpus of patient data and less studied mutations.

## VI. SUMMARY AND FUTURE WORK

**I**n this paper we examined a unique, largely unexplored corpus of clinical notes and a related collection of genetic mutation test results. We sought to gain insight into the hidden themes and framework of unstructured free-text medical notes, and to examine the parallels between that text and the genetic mutations of the patients it describes.

We began by accessing the clinical notes and genetic mutation records of several thousand patients from a private database of patient electronic health records assembled by Memorial Sloan-Kettering Cancer Center. These records were carefully prepared for analysis through several data pre-processing techniques in order to reduce noise and inconsistency which might lead to misleading or inconclusive results.

From our exploration of the clinical text notes we discovered large clusters of patients concerned with a variety of issues from the treatments and effects of specific cancers to broad interests in alternative medicine or opinions on surgical risk. We determine that there is a large amount of overlap between several of the main sections provided in typical clinical notes: Chief Complaint, History of Present Illness, Impressions and Plan, and Review of Systems, however other sections such as Social History and Family History contain unique details which are largely unrelated to the remainder of the note.

From an examination of the correlations between clinical note content and the available panel of mutation data, we successfully identified several genotype-phenotype relationships that have been independently identified through other methods of research. We consider this a successful proof-of-concept to motivate further refinement of our topic modeling approach.

Regarding future work, we are interested in the following directions. irst, we want to train topic models on sentences from documents and aggregate the resulting sentence representation per document to obtain a sharper estimate of topics and retain additional contextual detail. Next we wish to explore the entire series of patient notes over time to generate a temporal, naturally hierarchical topic representation of the clinical notes. Additionally, we wish to investigate a broader selection of mutation-topic associations with additional genetics data. Finally, we would like to explore other applications of topic models for clinical notes such as automated patient identification for clinical trials.

